# Genetic Analysis of Lodging in Diverse Maize Hybrids

**DOI:** 10.1101/185769

**Authors:** Sara J. Larsson, Jason A. Peiffer, Jode W. Edwards, Elhan S. Ersoz, Sherry Flint-Garcia, James B. Holland, Michael D. McMullen, Mitchell R. Tuinstra, M. Cinta Romay, Edward S. Buckler

## Abstract

Damage caused by lodging is a significant problem in corn production that results in estimated annual yield losses of 5-20%. Over the past 100 years, substantial maize breeding efforts have increased lodging resistance by artificial selection. However, less research has focused on understanding the genetic architecture underlying lodging. Lodging is a problematic trait to evaluate since it is greatly influenced by environmental factors such as wind, rain, and insect infestation, which make replication difficult. In this study over 1,723 diverse inbred maize genotypes were crossed to a common tester and evaluated in five environments over multiple years. Natural lodging due to severe weather conditions occurred in all five environments. By testing a large population of genetically diverse maize lines in multiple field environments, we detected significant correlations for this highly environmentally influenced trait across environments and with important agronomic traits such as yield and plant height. This study also permitted the mapping of quantitative trait loci (QTL) for lodging. Several QTL identified in this study overlapped with loci previously mapped for stalk strength in related maize inbred lines. QTL intervals mapped in this study also overlapped candidate genes implicated in the regulation of lignin and cellulose synthesis.

---

Lodging resistance is a critically important trait for maize hybrids in mechanized maize production systems. It is a major problem in corn production causing harvest difficulties and resulting in annual yield losses of 5 to 20% (Zuber and Kang, 1978; Flint-Garcia et al., 2003a). Lodging results from a plant’s inability to maintain an upright position when challenged with external factors including insect damage, fungal infestation, and weather conditions, such as rain and wind. Extensive breeding efforts have been made to develop lines with increased lodging resistance (Duvick, 2005). Yet, lodging remains an important criterion in maize improvement due to higher planting densities, higher soil fertility levels, and ever changing environmental factors. Often, lodging is classified into two categories – that occurring at the stalk and that occurring at the root (Lee and Tracy, 2009).

Stalk breakage can result in severely limited yield due to loss when breakage occurs below the primary ear or lack of photosynthetic surface area regardless of the location of stalk breakage (Ching et al., 2010). During a plant’s vegetative growth stage (V5-V8 and V12-R1), rapid growth of the internodes weakens cell walls, increasing the probability for the stalk to break when exposed to strong winds (Ching et al., 2010). When the plant has reached mature height, stalk lodging risks are moderate as lignin and other structural material strengthen the cell walls and stalk (Pedersen et al., 2005). Stalk lodging can also occur later in the season near harvest when the ear is fully developed and heavier and the stalk cannot support it. The weakness of the stalk later in the season is often confounded by insect infestations (e.g., by European corn borer, *Ostrinia nubilalis Hubner*) and stalk rots.

Root lodging is most often evaluated as the proportion of lodged plants per plot at maturity. A plant is considered to be root lodged when it is tilting >30° (e.g. Bruce et al., 2001; Landi et al., 2007). Root lodging in maize is affected by root characteristics including the number of roots on upper internodes, total root volume, root angle from vertical, and diameter of roots (Ennos, 1993). With a weakened root system, the plant is prone to wind damage resulting in snapping or buckling of the stalk at the base of the plant, or roots being pulled out of the soil. The risk for root lodging is highest during the mid-vegetative stage before brace roots are fully developed (Ennos, 1993). Root lodging early in the season is rarely devastating since a plant can regain an upright growing pattern due to its plasticity within a week with no negative effect on yield. This is not the case after the plant has fully matured (Zhang et al., 2011).

Accurate evaluations of genotypes for stalk and root lodging are difficult because of the influences of environmental factors that are not easy to control or replicate. To evaluate both stalk and root lodging under controlled wind conditions, DuPont Pioneer has developed a mobile wind machine that can generate winds up to 100 mph (Barreiro et al., 2008). Several other indirect methods can be used to evaluate potential resistance to stalk lodging like stalk crushing (Zuber et al., 1980), rind puncture resistance (RPR)(Djordjevic and Ivanovic, 1996, Peiffer et al., 2013), stalk water content (Djordjevic and Ivanovic, 1996), or near infrared (NIR) analysis of stalk tissue (Hu et al., 2012). For root lodging, alternative ways to determine susceptibility are by vertical-pull resistance (Ennos, 1993), measure of root volume by water replacement, and recording the weight of the root clump (Jenison et al., 1981).

Selection for some of those indirect traits, like increased RPR, has reduced stalk lodging (Albrecht and Dudley, 1987; Dudley 1994; Abedon et al., 1999). However, there are disadvantages to selecting genotypes with increased rind thickness, as thicker rinds may divert limited carbohydrates from kernel fill. This has the potential to result in lower yields (Davis and Crane, 1976). Other studies have reported a negative correlation between increased stalk strength and grain yield (Martin and Russell, 1984; Rhen and Russell, 1986). However, Colbert et al. (1984) found a significant positive correlation, while still other studies have observed no correlation between increases in stalk strength and other morphological traits (Djordjevic and Ivanovic, 1996). This suggests the relationships observed among traits may be strongly dependent upon both the germplasm surveyed and the testing environment.

A number of studies have been performed to better understand the genetic architecture underlying lodging. Genetic loci implicated in stalk strength have also been genetically mapped. Overall, these findings suggest that stalk strength is a highly complex trait controlled by a large number of alleles, each with small effects, and effective loci are not necessarily shared among different populations (Flint-Garcia et al., 2003a; Flint-Garcia et al., 2003b; Ching et al., 2010; Hu et al., 2012; Peiffer et al., 2013). Few QTL have been mapped for root lodging. One of the QTL identified to control root lodging is the root-ABA1 QTL on chromosome 2 (Landi et al., 2007). Moreover, QTL have been mapped for a number of root traits correlated with root lodging (e.g. Hochholdinger and Tuberosa, 2009).

As a result of their role in cell rigidity, lignin and cellulose content have been shown to influence stalk strength (Pedersen et al., 2005) and root lodging (Zhang et al., 2011). Nonetheless, natural variation in lignin content often has little impact on these traits (Pedersen et al., 2005). Much of the previously noted influence of lignin and cellulose is due to the observation of large rare effects such as mutant alleles of the *brown midrib* loci (*bm1* and *bm3*) (Vignols et al., 1995). Genes underlying these loci encode cinnamyl alcohol dehydrogenease (CAD) and a caffeic O-methyl transferase (COMT) of the lignin synthesis pathway (Sattler et al., 2010). Similarly, twelve *CesA* genes in the cellulose pathway are involved in secondary cell wall formation and have also been found to influence stalk strength (Appenzeller et al., 2004).

In this study, we examined hybrids from crosses of recombinant inbred lines (RILs) of the maize nested association mapping population (NAM) (McMullen et al., 2009) to the male tester PHZ51. These hybrids were grown in eight different field environments and their progenitors were genotyped (Buckler et al., 2009; McMullen et al., 2009). Five of the field locations were exposed to naturally occurring lodging events and each possessed substantial variation for lodging damage among genotypes. To relate this lodging variation to genetic diversity, we employed joint-linkage mapping of family-nested QTL. Several QTL were identified for lodging including some that overlapped with previously identified loci for lodging and related traits. However, several new QTL were also discovered.

## MATERIALS AND METHODS

### Germplasm

In this study we used hybrids of recombinant inbred lines (RILs) of the maize NAM population (McMullen et al., 2009) crossed to the male tester PHZ51. NAM was created by selecting 25 inbreds to maximize diversity, and crossing them to the reference inbred, B73. From each of the 25 families, 200 progeny were chosen, self-pollinated for five generations and subsequently sib-mated. This resulted in a mapping population of about 5,000 RILs. From these RILs, we selected a subset of 60-70 lines in each family (with the exception of the popcorn family, Hp301, which was wholly omitted) and created hybrids by crossing to the male tester PHZ51, a non-stiff stalk line developed by DuPont Pioneer with expired Plant Variety Protection (ex-PVP). The selection of NAM female inbreds for hybrid development was based on flowering time (Buckler et al., 2009). The earliest RILs from late families and the latest RILs from early families were selected to reduce flowering time variation and make hybrid production by isolation plots manageable.

### Phenotypic Evaluation

Hybrids were evaluated at Sandhills NC, Columbia MO, West Lafayette IN, and Slater IA in the summer of 2010, as well as Kinston NC, Columbia MO, West Lafayette IN, and Ames IA in the summer of 2011. All environments were cultivated in a conventional manner with respect to fertilization, weed, and pest management. Hybrids were planted in two-row plots with a single replication per environment. The experiment was blocked by family to avoid competition for space and light interception resulting from a lack of uniform height variation. Hybrids were randomized within blocks, and blocks were randomized within each environment. The hybrid B73xPHZ51 was used as a common check across all families and environments. In addition, the non-B73 parent crossed to PHZ51 was used as a second check within each family.

All phenotypic data were collected on a plot basis. Days from planting until half the plants in a plot shed pollen or had a visible silk was used as the criterion to measure days to anthesis and days to silk, respectively. All other traits were measured after flowering or at full maturity. Data were collected for height, leaf dimensions, and node counts after flowering when the plants had reached their full development. Plant height was measured as the distance from the soil line to the ligule of the flag leaf, and ear height as the distance from the soil line to the node of the primary ear. Leaf length and width were measured as the maximum length and width of the leaf below the primary ear. Numbers of nodes were separated into the number of nodes from the soil line to the node of the primary ear and the number from the node above the primary ear to the tassel. Root lodging was determined as the fraction of lodged plants within a plot. A plant was designated as root lodged when it leaned 30 degree or greater from vertical. Stalk lodging was measured as the proportion of plants in a plot with a broken stalk at or below the primary ear. Yield was measured using a two-row combine and moisture was recorded at the time of harvest. Yield was then adjusted to 15.5% moisture content and expressed in tons per hectare.

Due to weather conditions, five naturally occurring lodging events were examined in five unique field environments. In 2010, Columbia, MO and Slater, IA environments underwent lodging at flowering. That same year, the Sandhills, NC environment lodged after flowering. In 2011, the Columbia, MO environment underwent lodging early in the season before flowering and West Lafayette, IN lodged late in the season after flowering.

### Genotypic Evaluation

Genotypic data for joint linkage mapping of family-nested QTL was collected as previously described (McMullen et al., 2009; Buckler et al., 2009). In total, 1,106 markers were scored on an Illumina GoldenGate Assay across the NAM RILs. Among these markers, missing genotype calls were imputed as the weighted average of the flanking markers. Weights were derived from the missing marker’s genetic distance to each adjacent marker as previously described (McMullen et al., 2009; Buckler et al., 2009).

### Statistical Analysis

Statistical analyses were performed using SAS (SAS Institute, 2004) software, and R (R Development Core Team, 2008) scripts. Best linear unbiased estimates (BLUEs) for the genotype values of all phenotypes across and within environments were calculated using the LSMEANS statement in PROC MIXED of SAS software while fitting genotypes as fixed effects and accounting for environment, block within environment, range position, and row position as random effects. Heritabilities of the lodging traits for the five environments were calculated following Hung et al. (2011).

After calculating BLUE genotype values across and within environments, the base package of R was used to calculate Pearson correlation coefficients and to study relationships between phenotypes at the genotypic level across and within environments and NAM families. In the statistical analyses detailing lodging conditional on flowering stage, phenotypes from Columbia, MO and Slater, IA in 2010 were grouped to define lodging occurring at flowering. Using the same principal, data from Sandhills, NC in 2010 and West Lafayette, IN in 2011 were grouped to detail lodging occurring after flowering.

To better characterize genetic architecture, joint linkage mapping of family-nested QTL explaining variation of the BLUE genotype values across and within environments was performed using PROC GLMSELECT in SAS. The model included the “family” term in order to explain variation between NAM families, and the set of 1,106 markers were nested within each NAM family. These markers were then selected for model inclusion based on their covariation with BLUE genotype values for each phenotype across and within environments in a stepwise manner (Buckler et al., 2009). For the stepwise model selection procedure, inclusion and exclusion of family nested markers were discerned by comparison with a null distribution based on permutation testing. The *p*-value derived from the null distribution for model inclusion was 0.001 at an alpha level of 0.05. Joint linkage mapping of family-nested QTL was performed both with and without accounting for covariation of flowering time (as measured by days to anthesis) within the model.

## RESULTS

### Phenotypic evaluation

All five environments experienced substantial lodging and variation for root and stalk lodging (Table 1). About 85-99 percent of plots in the five environments had one or more lodged plants. In the MO11 environment, which was damaged by a storm early in the season, the majority of the lodging was stalk lodging. The same pattern was observed in the NC10 environment; however, IA10 contrasted this trend and had a high proportion of root lodging and low proportion of stalk lodging. Relative to these environments, MO10 and IN11 had high percentages of both root lodging and stalk lodging.

**Table 1.**
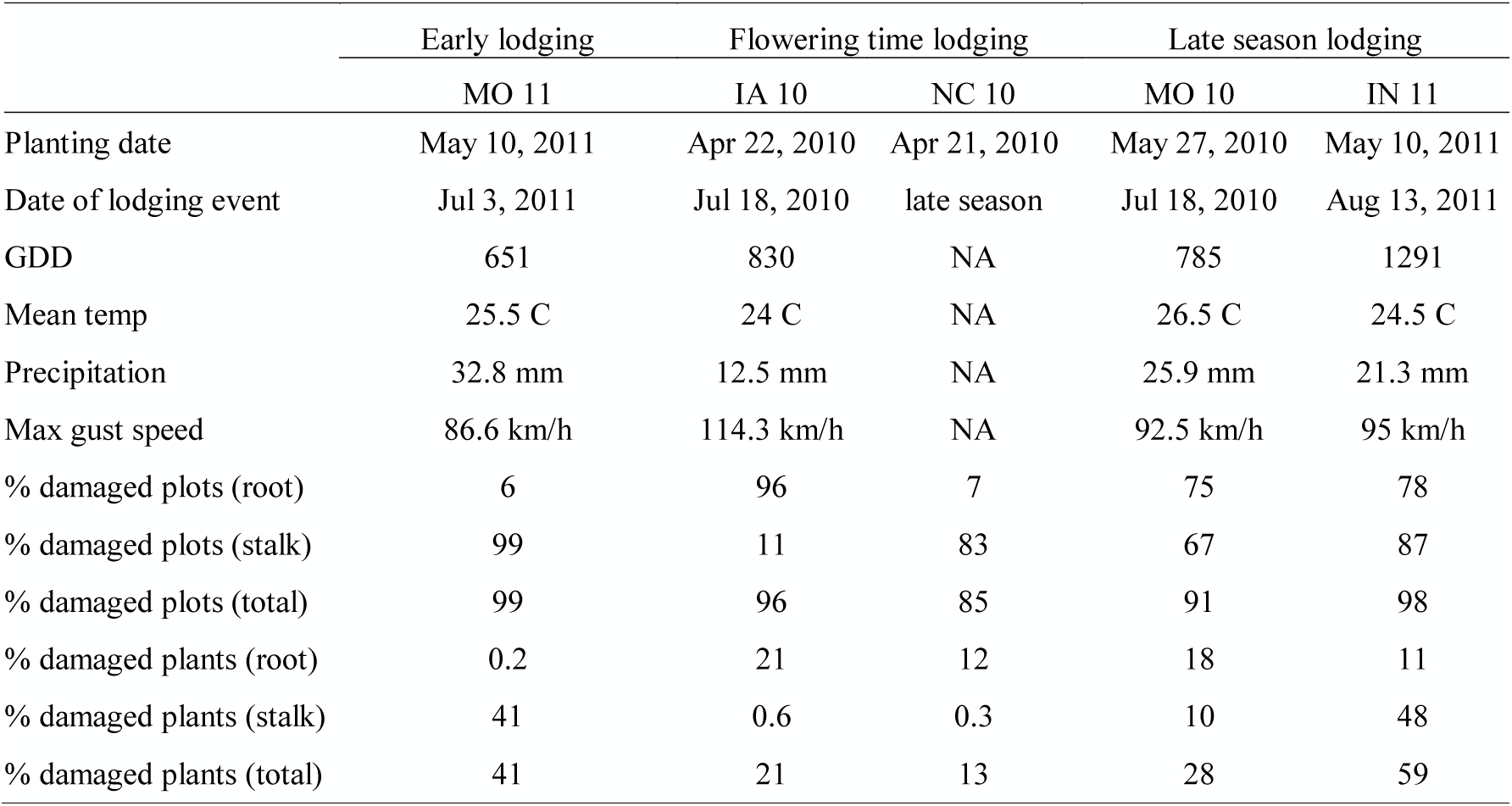
Date of planting and storm events, and information on weather conditions at the day of the event (Steremberg 2012), percent of plots per environment damaged by lodging. Growing degree days (GDD) are calculated with a base temperature of 10 C until the day of the event.

|  | Early lodging | Flowering time lodging |  | Late season lodging |  |
| --- | --- | --- | --- | --- | --- |
|  | MO 11 | IA 10 | NC 10 | MO 10 | IN 11 |
| Planting date | May 10, 2011 | Apr 22, 2010 | Apr 21, 2010 | May 27, 2010 | May 10, 2011 |
| Date of lodging event | Jul 3, 2011 | Jul 18, 2010 | late season | Jul 18, 2010 | Aug 13, 2011 |
| GDD | 651 | 830 | NA | 785 | 1291 |
| Mean temp | 25.5 C | 24 C | NA | 26.5 C | 24.5 C |
| Precipitation | 32.8 mm | 12.5 mm | NA | 25.9 mm | 21.3 mm |
| Max gust speed | 86.6 km/h | 114.3 km/h | NA | 92.5 km/h | 95 km/h |
| % damaged plots (root) | 6 | 96 | 7 | 75 | 78 |
| % damaged plots (stalk) | 99 | 11 | 83 | 67 | 87 |
| % damaged plots (total) | 99 | 96 | 85 | 91 | 98 |
| % damaged plants (root) | 0.2 | 21 | 12 | 18 | 11 |
| % damaged plants (stalk) | 41 | 0.6 | 0.3 | 10 | 48 |
| % damaged plants (total) | 41 | 21 | 13 | 28 | 59 |

Heritabilities of the traits were moderate. Heritabilities on a plot bases ranged from 0.12 for stalk lodging to 0.24 for root lodging, and mean heritabilities ranged from 0.23 for stalk lodging to 0.41 for root lodging (Table 2).

**Table 2.** Heritability (H^2^) on a plot and mean basis for root lodging, stalk lodging, and total lodging.

| | $H^2$ Plot | $H^2$ Mean |
| --- | --- | --- |
| Root lodging | 0.24 | 0.41 |
| Stalk lodging | 0.12 | 0.23 |
| Total lodging | 0.16 | 0.30 |

### Trait correlations

Both root and stalk lodging are highly influenced by environmental factors such as weather conditions, especially wind and water. Correlations of root lodging among the five locations were strongest –0.31 – between environments IA10 and MO10 (Table 3). The IN11 environment shows correlation with MO10 and IA10. The NC10 environment, which was exposed to late season lodging, had a negative correlation with root lodging in all other environments. Overall NC10 did not possess much root lodging, since the probability of root lodging is higher earlier in the growing season. For stalk lodging, the strongest correlations were between IA10 and MO10 (0.17), NC10 and IN11 (0.25), as well as MO11 and IN11 (0.21), each of which were statistically significant (*p*-value = <0.0001). For total lodging the two highest correlations were between MO10 and IA10, and MO11 and IN11. MO10 and IA10 were both exposed to severe weather conditions on the same date and stage of development.

**Table 3.** Correlation between root lodging (A), stalk lodging (B), and total lodging (C) across the five environments. Upper right half of the tables reports r-values and lower left half reports *p*-values.

| A |  |  |  |  |  |
| --- | --- | --- | --- | --- | --- |
|  | Root MO 10 | Root IA 10 | Root NC 10 | Root IN 11 | Root MO 11 |
| Root MO 10 (middle) | ----- | 0.31 | -0.02 | 0.18 | 0.02 |
| Root IA 10 (middle) | <0.0001 | ----- | -0.04 | 0.23 | 0.00 |
| Root NC 10 (late) | 0.5802 | 0.2170 | ----- | -0.06 | -0.01 |
| Root IN 11 (late) | <0.0001 | <0.0001 | 0.0542 | ----- | 0.05 |
| Root MO 11 (early) | 0.4893 | 0.9560 | 0.8674 | 0.0598 | ----- |
| B |  |  |  |  |  |
|  | Stalk MO 10 | Stalk IA 10 | Stalk NC 10 | Stalk IN 11 | Stalk MO 11 |
| Stalk MO 10 (middle) | ----- | 0.17 | 0.00 | 0.03 | -0.05 |
| Stalk IA 10 (middle) | <0.0001 | ----- | -0.07 | 0.02 | -0.02 |
| Stalk NC 10 (late) | 0.9615 | 0.3410 | ----- | 0.25 | 0.07 |
| Stalk IN 11 (late) | 0.2950 | 0.6552 | <0.0001 | ----- | 0.21 |
| Stalk MO 11 (early) | 0.1303 | 0.4865 | 0.017 | <0.0001 | ----- |
| C |  |  |  |  |  |
|  | Total MO 10 | Total IA 10 | Total NC 10 | Total IN 11 | Total MO 11 |
| Total MO 10 (middle) | ----- | 0.28 | 0.08 | 0.16 | 0.00 |
| Total IA 10 (middle) | <0.0001 | ----- | 0.06 | 0.17 | 0.09 |
| Total NC 10 (late) | 0.0079 | 0.0720 | ----- | 0.20 | 0.08 |
| Total IN 11 (late) | <0.0001 | <0.0001 | <0.0001 | ----- | 0.21 |
| Total MO 11 (early) | 0.9394 | 0.0066 | 0.0073 | <0.0001 | ----- |

All environments were grouped by the time of lodging with respect to flowering time, i.e., if the lodging event occurred before, at, or after time of flowering. Environments in which lodging occurred at (MO10 and IA10) or after (NC10 and IN11) flowering showed the expected patterns of correlation. High correlations were observed between flowering traits (days to silk and days to anthesis) and plant and ear height. For the environments in which lodging occurred at flowering, significant (*p*-value = <0.0001) negative correlations between the traits and yield were observed. This was especially true of the lodging traits (Table 4). Negative correlations between stalk/total lodging and yield were also present in environments with lodging after flowering (Table 5). Within these environments, there was also a high correlation between lodging traits and plant and ear height.

**Table 4.** Correlations between the three lodging traits and other developmental traits measured in the middle environments, where lodging occurred at flowering. Upper right half of the table reports r-values and lower left half reports *p*-values. D2A = days to anthesis, D2S = days to silk.

|  | D2A | D2S | Ear Height | Plant Height | Leaf Length | Leaf Width | Root | Stalk | Total Lodging | Yield |
| --- | --- | --- | --- | --- | --- | --- | --- | --- | --- | --- |
| D2A | ----- | 0.90 | 0.07 | -0.15 | 0.11 | 0.03 | 0.18 | 0.06 | 0.18 | -0.13 |
| D2S | <0.0001 | ----- | 0.10 | -0.14 | 0.17 | 0.06 | 0.18 | 0.07 | 0.18 | -0.17 |
| Ear Height | 0.0167 | 0.0007 | ----- | 0.63 | 0.33 | 0.16 | 0.22 | 0.01 | 0.21 | -0.21 |
| Plant Height | <0.0001 | <0.0001 | <0.0001 | ----- | 0.28 | 0.07 | 0.18 | -0.08 | 0.15 | -0.14 |
| Leaf Length | 0.0003 | <0.0001 | <0.0001 | <0.0001 | ----- | 0.16 | 0.14 | 0.07 | 0.15 | -0.20 |
| Leaf Width | 0.3573 | 0.0721 | <0.0001 | 0.0301 | <0.0001 | ----- | 0.16 | 0.08 | 0.18 | -0.12 |
| Root | <0.0001 | <0.0001 | <0.0001 | <0.0001 | <0.0001 | <0.0001 | ----- | 0.06 | 0.90 | -0.23 |
| Stalk | 0.0402 | 0.0185 | 0.7344 | 0.0047 | 0.0361 | 0.0149 | 0.0344 | ----- | 0.43 | -0.21 |
| Total | <0.0001 | <0.0001 | <0.0001 | <0.0001 | <0.0001 | <0.0001 | <0.0001 | <0.0001 | ----- | -0.28 |
| Yield | <0.0001 | <0.0001 | <0.0001 | <0.0001 | <0.0001 | 0.0003 | <0.0001 | <0.0001 | <0.0001 | ----- |

**Table 5.** Correlations between the three lodging traits and other developmental traits measured in the late environments, where lodging occurred after flowering. Upper right half of the table reports r-values and lower left half reports *p*-values.

|  | D2A | D2S | Ear<br>Height | Plant<br>Height | Leaf<br>Length | Leaf<br>Width | Brace<br>Roots | Nodes<br>Ear-Tassel | Nodes<br>Roots-Ear | Root | Stalk | Total<br>Lodging | Yield |
| --- | --- | --- | --- | --- | --- | --- | --- | --- | --- | --- | --- | --- | --- |
| D2A | ----- | 0.73 | 0.10 | -0.06 | 0.22 | -0.02 | 0.04 | -0.01 | 0.17 | -0.09 | -0.11 | -0.16 | 0.05 |
| D2S | 0.1798 | ----- | 0.16 | -0.05 | 0.21 | 0.02 | 0.04 | -0.06 | 0.23 | -0.05 | -0.12 | -0.14 | 0.07 |
| Ear<br>Height | 0.1462 | <0.0001 | ----- | 0.75 | 0.15 | 0.09 | 0.01 | 0.10 | 0.50 | 0.29 | 0.40 | 0.50 | 0.09 |
| Plant<br>Height | 0.7157 | 0.0005 | <0.0001 | ----- | 0.05 | -0.04 | 0.05 | 0.31 | 0.40 | 0.41 | 0.43 | 0.60 | 0.15 |
| Leaf<br>Length | 0.0002 | <0.0001 | <0.0001 | <0.0001 | ----- | 0.09 | 0.10 | -0.02 | 0.09 | -0.16 | -0.04 | -0.13 | 0.08 |
| Leaf<br>Width | 0.4019 | 0.4810 | 0.5355 | 0.0004 | 0.0003 | ----- | -0.02 | -0.07 | 0.02 | -0.13 | -0.05 | -0.12 | 0.09 |
| Brace<br>Roots | 0.0034 | 0.8139 | 0.0404 | 0.0002 | 0.4572 | 0.0045 | ----- | 0.08 | 0.20 | -0.08 | -0.04 | -0.08 | 0.04 |
| Nodes<br>Ear-Tassel | <0.0001 | <0.0001 | <0.0001 | <0.0001 | 0.0007 | 0.4145 | 0.3058 | ----- | -0.03 | 0.20 | 0.08 | 0.19 | 0.00 |
| Nodes<br>Roots-Ear | 0.0645 | 0.0432 | 0.0788 | <0.0001 | 0.0354 | 0.0941 | <0.0001 | <0.0001 | ----- | 0.18 | 0.21 | 0.28 | 0.05 |
| Root | 0.0029 | 0.0017 | 0.0838 | <0.0001 | <0.0001 | <0.0001 | <0.0001 | <0.0001 | <0.0001 | ----- | -0.05 | 0.51 | 0.03 |
| Stalk | 0.1348 | 0.0001 | <0.0001 | <0.0001 | 0.1433 | 0.0365 | 0.0019 | <0.0001 | <0.0001 | 0.0675 | ----- | 0.83 | -0.19 |
| Total | 0.0039 | <0.0001 | <0.0001 | <0.0001 | <0.0001 | <0.0001 | <0.0001 | <0.0001 | <0.0001 | <0.0001 | <0.0001 | ----- | -0.15 |
| Yield | 0.1887 | 0.1427 | 0.0331 | 0.0031 | 0.0089 | 0.0017 | 0.8719 | 0.1047 | <0.0001 | 0.3072 | <0.0001 | <0.0001 | ----- |

Genotypes within each environment were grouped by percentage (in increments of 10%) of lodging damage. Larger proportions of damaged plants within a group resulted in lower average yields (Table 6). However, genotypes with both low and high levels of resistance to lodging have the potential for high yields in good season environments, without lodging events (data not shown). Over all environments, the correlation between percentage of lodging and yield was −0.10 with a *p*-value of 0.0009.

**Table 6.** Average yield in T/ha of genotypes grouped according to percentage of total lodging damage per plot for individual environments.

| Percent total lodging | MO 10 | IA 10 | NC 10 |
| --- | --- | --- | --- |
| 0 - 10 | 6.65 | 6.72 | 4.82 |
| 11 - 20 | 6.01 | 6.50 | 4.51 |
| 21 - 30 | 5.34 | 5.96 | 4.27 |
| 31 - 40 | 4.04 | 6.07 | 3.73 |
| 41 - 50 | 1.30 | 5.90 | 3.57 |
| 51 - 60 | 1.12 | 4.48 | 3.13 |
| 61 - 70 | 0.00 | 1.96 | 1.81 |
| 71 - 80 | 0.00 | NA | 2.22 |
| 81 - 90 | 0.00 | 1.86 | 1.79 |
| 91 - 100 | 0.00 | 0.00 | 0.00 |

### Joint linkage mapping

We performed joint linkage mapping of family-nested QTL for root lodging, stalk lodging, and total lodging. Analyses were done for the grouped environments, lodging at and after flowering, as well as for each environment independently. Most family-nested QTL that mapped within a single environment were shared across the grouped environments. For the two grouped environments, four family-nested QTL for stalk lodging, ten family-nested QTL for root lodging, and nine family-nested QTL for total lodging were identified (Figure 1, Table 7, Supplemental Table 1). Joint linkage mapping was performed both excluding and including flowering time as a covariate to account for stage of maturity at the time of the lodging event. Including flowering time in the model did not have a significant effect on mapping results (data not shown).

**Figure 1.**
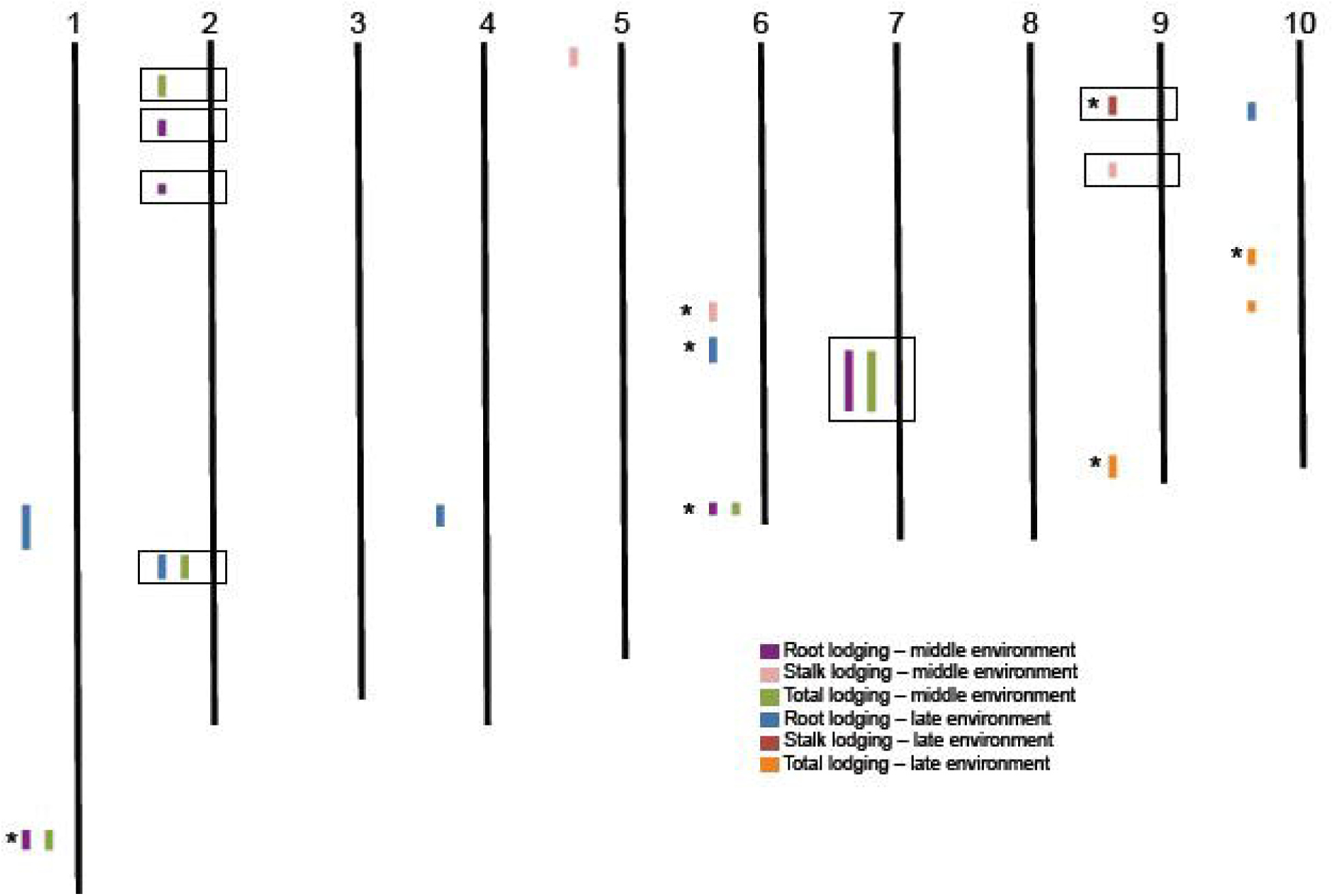
Distribution of QTL mapped using joint linkage mapping across the ten chromosomes. * indicates QTL intervals overlapping with results from stalk strength study in the NAM inbred population. Boxes indicate QTL overlapping with candidate genes.

**Table 7.** Mapped QTL and overlapping intervals with other lodging studies.

|  | Chr | cM start | cM end | Peiffer et al. 2013 | Ching et al. 2010 | Flint-Garcia et al. 2003a | Flint-Garcia et al. 2003b | Hu et al. 2012 |
| --- | --- | --- | --- | --- | --- | --- | --- | --- |
| Root (late) | 1 | 94.9 | 96.5 |  |  | X |  |  |
| Root / Total (middle) | 1 | 175.7 | 176.9 | X | X |  |  | X |
| Total (middle) | 2 | 41.5 | 41.8 | X |  |  |  |  |
| Root (middle) | 2 | 54.9 | 56.8 |  |  |  |  |  |
| Root (middle) | 2 | 72.1 | 72.1 |  |  |  |  | X |
| Root (late) / Total (middle) | 2 | 97.5 | 98.9 | X |  | X |  |  |
| Root (late) | 4 | 93.2 | 93.2 |  |  |  |  |  |
| Stalk (middle) | 5 | 10.1 | 10.9 |  |  |  |  |  |
| Stalk (middle) | 6 | 41.5 | 42.1 | X |  |  |  |  |
| Root (late) | 6 | 43.4 | 43.7 | X |  |  |  |  |
| Root / Total (middle) | 6 | 101 | 106.6 | X |  |  | X | X |
| Root / Total (middle) | 7 | 69.8 | 69.8 |  |  |  |  |  |
| Stalk (late) | 9 | 44.5 | 45.2 |  |  |  |  |  |
| Stalk (middle) | 9 | 47 | 47.2 |  |  |  |  |  |
| Total (late) | 9 | 89.8 | 93.5 |  |  |  | X |  |
| Root (late) | 10 | 40.1 | 40.6 | X |  |  | X |  |
| Total (late) | 10 | 43.4 | 43.8 |  |  |  | X |  |
| Total (late) | 10 | 44.7 | 44.8 |  |  |  | X |  |

## DISCUSSION

Negative correlations between lodging traits and yield were observed in testcross hybrids of the NAM population evaluated in eight environments, five of them with natural lodging events (Table 4 and 5). It has been suggested that there is a tradeoff between breeding for higher yield and more lodging resistant lines. It has been a concern for many years that breeding for lodging resistance by increasing stalk strength decreases yield (Rhen and Russell, 1986). This reasoning is based on the sink-source relationship of available carbon in the plant. If more carbon is expended to strengthen stalks, there is less available for grain fill later in the season (Davis and Crane, 1976).

In environments with late season lodging, correlations between plant and ear height and lodging traits were observed (Table 5). Similar relationships between height and lodging have been reported in previous studies (e.g. Flint-Garcia et al., 2003a, 2003b; Holthaus and Lamkey, 1995). Higher ear placement and heavier ears in the late season in combination with weaker roots or stalks will more likely result in lodging, compared to shorter plants with lower ear placement. In addition, there is a relationship between total plant height and yield (Duvick, 2005). Plants increase in biomass and photosynthetic rates and fix more carbon that can be allocated to the ear as grain yield until reaching an optimal height. As such, it is not as simple as exclusively breeding for shorter genotypes to avoid lodging when seeking to increase grain yield.

Family-nested QTL for root, stalk, and total lodging were identified (Supplemental Table 1). Fewer family-nested QTL were identified than previously mapped for stalk strength using RPR in the full set of NAM inbreds (Peiffer et al., 2013). The inbred study identified over 18 family-nested QTL. One explanation is the difference in population size. The current study only used about one-third of the lines compared to the 5,000 RIL in the NAM population. Second, the traits in the current study were caused by environmental condition — proportion of plots damaged by weather—which is more difficult to evaluate and replicate, in comparison to the force it takes to penetrate the stalk reported in Peiffer et al. (2013). Overall, this suggests that the traits are controlled by a large number of loci with small effects. It is likely that we were only able to identify family-nested QTL with the larger effects. However, we believe our results to be robust.

Seven of the family-nested QTL mapped in this study are located in the same marker intervals on the NAM map as family-nested QTL identified using RPR (Peiffer et al., 2013). A large number of studies of lodging and stalk strength have been performed. We compared our results with these studies to assess overlap of the different phenotyping strategies. Peiffer et al. (2013) measured stalk strength in maize using RPR in the inbred NAM population used as females in hybrid development for this study. Flint-Garcia et al. (2003a, 2003b) also studied stalk strength in maize using RPR across multiple environments and multiple populations selected for high and low stalk strength. Ching et al. (2010) measured stalk strength using mechanical force, and Hu et al. (2012) used both RPR and NIR. Overall, our results show considerable overlap with these studies for stalk lodging (Table 7) (Flint-Garcia et al., 2003a, 2003b; Ching et al., 2010; Hu et al., 2012). In addition, the family-nested QTL for root lodging located on chromosome 2 (97.2 – 98.9 cM) mapped within the same virtual bin, 2.04, as the root-ABA QTL that was previously mapped and is known to influence root lodging (Landi et al., 2007).

A list of 50 genes with known involvement in lignin synthesis, phenylpropanoid pathway, vegetative phase change and cellulose was compiled (Supplemental Table 2). This list was compared to mapped family-nested QTL in this study. Ten of the fifty genes are located within 3 Mb of the peak marker for a lodging QTL. Four of the genes are involved in the lignin synthesis, one in the phenylpropanoid pathway, two are involved in the vegetative phase change, and three genes are involved in the cellulose synthesis pathway.

Here we have performed one of the largest public studies detailing the genetic architecture of natural lodging in diverse hybrids. This study was performed across five unique environments, each differently affecting stalk and root lodging. The diverse populations and rapid LD decay have permitted mapping of QTL for root, stalk, and total lodging. Several identified QTL overlapped with previous studies on stalk strength. In addition, candidate genes influencing stalk and root composition were located within the intervals of the mapped family-nested QTL. This study has provided a deeper understanding of natural lodging in maize.

## ACKNOWLEDGEMENTS

This work was funded by NSF Plant Genome Program (IOS 0820619 and 1238014) and USDA-ARS. Graduate work of SJL, work of ESE, and IA10 trials were partially funded by Syngenta.

